# Hormetic Shifting of Redox Environment by Pro-Oxidative Resveratrol Protects Cells Against Stress

**DOI:** 10.1101/045567

**Authors:** Annabell Plauth, Anne Geikowski, Susanne Cichon, Silvia J. Wowro, Linda Liedgens, Morten Rousseau, Christopher Weidner, Luise Fuhr, Magdalena Kliem, Gail Jenkins, Silvina Lotito, Linda J. Wainwright, Sascha Sauer

## Abstract

Resveratrol has gained tremendous interest owing to multiple reported health-beneficial effects. However, the underlying key mechanism of action of resveratrol remained largely controversial. Here, we demonstrate that under physiologically relevant conditions major biological effects of resveratrol can be attributed to the generation of oxidation products such as reactive oxygen species (ROS). At low hormetic concentrations (< 50 μM), treatment with resveratrol increased cell viability in a set of representative cell models, whereas application of quenchers of ROS completely truncated these beneficial effects. Notably, application of resveratrol led to mild, Nrf2-specific cellular gene expression reprogramming. For example, in primary human epidermal keratinocytes this resulted in a 1.3-fold increase of endogenous metabolites such as gluthathione (GSH) and subsequently in a quantitative reduction of the cellular redox environment by 2.61 mV mmol GSH (g protein)^-1^. In particular in resveratrol pre-treated cells, after external application of oxidative stress by using 0.8 % ethanol, endogenous ROS generation was consequently reduced by 24 %. In contrast to the common perception that resveratrol acts mainly as a chemical antioxidant or as a target protein-specific ligand, we propose that effects from resveratrol treatment are essentially based on oxidative triggering of cells. In relevant physiological microenvironments this effect can lead to hormetic shifting of cellular defence towards a more reductive state to improve resilience to oxidative stress in a manner that can be exactly defined by the redox-environment of the cell.

## Introduction

Polyphenols represent a large collection of natural products featuring health-beneficial effects^1^. Resveratrol (3,5,4'-trihydroxy-trans-stilbene, RSV), an antimicrobial phytoalexin originally found in white hellebore (*Veratrum grandiflorum O Loes*) and later in red grapes and other plants, is one of the most prominent polyphenols. Early studies indicated cancer chemo-preventive properties of RSV ^2^. Over the last 15 years, numerous studies claimed additional benefits including cardioprotective and anti-aging effects ^3^. Consequently, a number of products based on RSV have been developed for dietary and dermatological application ^4,5^. Nevertheless, the efficiencies of RSV treatments and underlying mechanisms of action remained largely controversial. For example, RSV had been suggested to modulate estrogen receptor activity ^6^, or act as a caloric mimetic by directly increasing the enzymatic activity of the histone deacetylase sirtuin 1 (SIRT1) ^7^. Recently, it was shown that inhibition of phosphodiesterase 4 (PDE4) by RSV increased intracellular amounts of the hunger signalling molecule cAMP ^8^. Notably, the reported interaction of RSV with these and further target proteins were in many cases low and unspecific (mostly in the mid micromolar range). In general, most of these studies assumed a proportional dose-response relationship of compounds, i.e. a conventional pharmacological (linear) threshold model ^9^

However, in contrast to the standard pharmacological model, hundreds of studies reported (unconsciously) beneficial effects of RSV at “low” but detrimental outcomes at “high” doses. Nevertheless, this potentially counterintuitive bi-phasic property of RSV was widely ignored ^10^. The large body of these data would hint to hormesis, a dose-response relationship that is characterized by low-dose stimulation and highdose inhibition, consistent with the Arndt-Schulz law, Hueppe’s rule and other terms describing a beneficial stimulation (of poisons) at low doses ^11, 12^. General acceptance of the hormesis concept for therapeutic application seems to remain low, due to the generally low stimulatory effects and particularly due to an often lacking mechanistic explanation of the underlying mode of action of so-called hormetic compounds.

Interestingly, polyphenols including RSV are considered as antioxidants. But depending on the chemical context RSV and other polyphenols can also become pro-oxidative^1^, a fact that is nevertheless often ignored. Depending on the reaction conditions resveratrol can be (auto-) oxidized to generate semiquinones and the relatively stable 4'-phenoxyl radical, which can produce reactive oxygen species (ROS) ^13, 14^. (Auto-) oxidative reactions of polyphenols are influenced by changing pH, particularly the presence of hydroxyl anions or organic bases ^15, 16^. Additionally, metal ions (e.g. iron II ions) facilitate oxidative reactions and further radical generation via Fenton reactions ^17^

This study aimed to connect fragmented pieces of the chemical and resulting biological properties of RSV to provide a conceptually comprehensive mechanistic understanding of the varying purported health-beneficial effects of RSV.

## Results and Discussion

### RSV is unstable under physiologically relevant conditions

The vast majority of studies seem to assume specific RSV-target protein interactions, which implies that RSV remains intact during treatments. However, after incubation in various media containing physiological concentrations of sodium bicarbonate (NaHCO_3_), a key component of water as well as buffer of blood and biological cells, RSV reacts efficiently, as indicated by striking yellowish colour changes (Fig. S1a and b). Light absorbance at characteristic RSV maximum (308 nm) decreased rapidly in water and cell culture media, both containing sodium bicarbonate (Fig. S1c and d, Table S1). After 16 hours incubation the absorption maximum of RSV was almost completely diminished. Furthermore, using a commonly applied fluorescence-based SIRT1 assay no enzymatic activation could be detected (Fig. S1e).

Oxidation of resveratrol at atmospheric oxygen level (21% O_2_, as usually applied in cell culture) ^16, 18^ could potentially be considered as non-physiological (in blood vessels the oxygen amount is roughly 14% and in tissues or tumours even lower (~1% O_2_)) ^19^. Here we show that the auto-oxidation of RSV is highly dependent on the presence of sodium bicarbonate and pH of the solvent ^14^ (Fig. S2 and S3), whereas decreasing oxygen partial pressure seems to have a comparably minor influence on the oxidation efficiency of RSV (Fig.S3a and b). These data suggest that oxidation of RSV can even take place in hypoxic microenvironments.

These results corroborate widely ignored findings that the stability and oxidation of RSV in physiologically relevant media is strongly influenced by pH and in particular the availability of hydroxyl anions ^16^,18,^20^. Interestingly, RSV reacted also efficiently in tap water (Fig. S4e left), which might explain further the often-reported perplexing low bioavailability of RSV *in vivo* ^21-23^. Although RSV could potentially be protected from protein carriers such as serum albumin ^24^, the entirety of these data makes it difficult to understand how RSV could exert compound-protein specific effects. These results further indicate that potential metabolisation of RSV, for example by oxidation of the enzymes of the CYP1 family, might play a minor role in physiological context, consistent with the usually extremely low amounts of detected metabolites of resveratrol ^21-23^.

### RSV produces ROS under physiologically relevant conditions

We next asked how the oxidising RSV could induce relevant biological effects. Notably, treatment of cells with RSV resulted in time‐ and concentration-dependent generation of intracellular ROS (Figure 1a and b and Fig. S4a).

**Figure 1.**
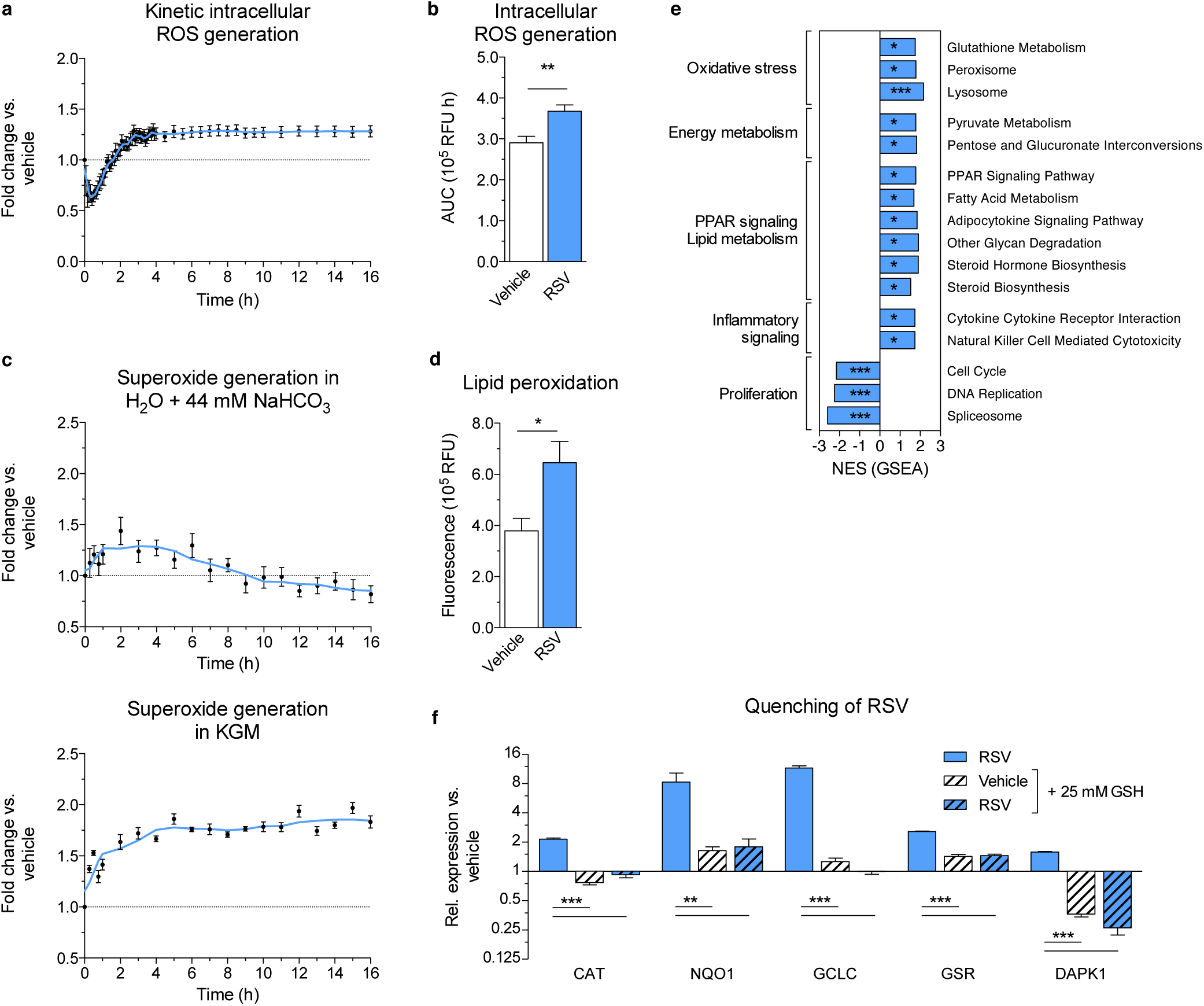
RSV produces ROS and triggers cellular defence. **(a and b)** Kinetic **(a)** and summed **(b)** intracellular intracellular ROS generation in primary NHEK cells after administration of 50 μM RSV. Values are mean ± s.e.m. (n = 6); AUC, area under the curve; RFU, relative fluorescence units. **(c)** Generation of superoxide in H_2_O and keratinocyte growth medium (KGM) after addition of 50 μM RSV for indicated time periods. Values are mean ± s.e.m. (n = 4). **(d)** RSV (50 μM) induced lipid peroxidation in NHEKs. Values are mean ± s.e.m. (n = 4); * *P* < 0.05 versus vehicle. **(e)** Selection of enriched KEGG pathways in NHEKs treated with 50 μM RSV using gene set enrichment analysis (GSEA). Values are normalized enrichment scores (NES, n = 4 for vehicle, n = 3 for RSV); *P < 0.05, **P < 0.01, *** *P* ≤ 0.001 versus vehicle. See also Table S2. **(f)** GSH considerably quenched RSV effects on gene expression. Values are mean ± s.e.m. (n = 4); ** *P* < 0.01; *** *P* ≤ 0.001 one-way ANOVA versus RSV.

Owing to the complexity of biological matrices it remains technically impossible to detect exactly unstable intermediate ROS. However, using defined solvents or a cell-free environment, we detected significantly increased amounts of ROS, including superoxide anions (Figure 1c), hydroxyl radicals (Fig. S4b-d) and hydrogen peroxide (Fig. S4e) ^14, 16^. For example, between 5.0 and 12.5 μM hydrogen peroxide (Fig. S4e) was generated depending on the concentration of RSV and sodium bicarbonate (Fig. S4e left), whereas (metabolic) scavengers such as pyruvate strongly depleted ROS (Fig. S4e right). As revealed by an antioxidant assay using trolox as control compound, in presence of sodium bicarbonate the anti-oxidative feature of ROS-producing RSV was strongly diminished (Fig. S4f left). In contrast, RSV showed roughly 2-fold higher anti-oxidative capacity in solvents lacking sodium bicarbonate (Fig. S4f right). These data suggest that RSV loses in part its anti-oxidative properties and becomes more pro-oxidative in physiological media containing sodium bicarbonate.

These data indicate that oxidation of RSV takes place under physiologically relevant conditions that differ significantly from experimental setups applied to analyse RSV by common bicarbonate-free enzymatic assays, or bicarbonate-free crystallization procedures that are mostly used for x-ray analyses. Notably, low concentrations of oxidation products of RSV such as ROS can mildly affect cellular biomolecules such as proteins and lipids (Figure 1d).

### Effects on primary human keratinocytes

We next asked how oxidation of RSV might influence physiological effects. Given the here observed oxidative effects in particular topological application of RSV seems a medially relevant approach, as is evident from the number of available dermatological products based on RSV. We thus focused in this study on potential protection of the human epidermis. Notably, due to ethical considerations and law, for physiological testing dermatological research applies *ex vivo* models such as the here used primary human keratinocytes that form the outer layer of the skin. Keratinocytes are known to build a tight layer of cells that can be used as epidermal grafts (Fig. S4g) ^25^, and these cells represent a prime target for lotions and emollients based on RSV ^26, 27^.

Firstly, we investigated potential oxidation of cellular components owing to oxidation products of RSV. Sixteen hours of treatment with 50 μM RSV slightly elevated lipid peroxidation in human primary keratinocyte (NHEK) cells, indicating mildly increased oxidation of cellular biomolecules (Figure 1d).

Moreover, to globally monitor cellular response to oxidation products of RSV, genome-wide RNA expression analyses revealed a slight but significantly increased expression of molecular pathways covering oxidative stress response and inflammatory signalling, as well as fatty acid metabolism (Figure 1e and Table S2). In contrast, processes linked to proliferation, DNA replication and cell cycle were down-regulated. Importantly, quenching of oxidation by adding strong reducing molecules such as 25 mM synthetic GSH significantly reversed expression of cell response marker genes (Figure 1f). As shown in a control experiment, quenching by synthetic GSH in reduced the levels of ROS derived from RSV in cell-free environment (Fig. S4g) and even more importantly within cells (Fig. S4i), suggesting that ROS derived from RSV mainly cause the observed gene expression effects. As tested with a small panel of unrelated cell models, despite cell-specific defence mechanisms the here observed gene expression events seem to some degree be independent from cellular background (Fig S5a-d, Table S3).

These data indicate to our knowledge for the first time that major gene expression events induced by RSV can be explicitly attributed to the development of oxidation products of RSV such as ROS, since depletion by molecular quenchers strongly truncated cellular response. The effects of RSV analysed here in mammalian cells might also underlie phytoalexin-based protection of plants against microbial infection^28^.

### Oxidative products of RSV cause hormetic effects

In a next step, we asked how oxidation products derived from RSV could potentially influence viability of cells. Using common cell viability assays, we observed increased cellular fitness up to about 50 μM RSV in treated NHEKs, whereas higher concentrations tend to produce toxic effects, leading to a typical bi-phasic, hormetic dose-viability curve (Figure 2a and b). Notably, in additional cellular models for fibroblasts and liver we observed similar bi-phasic dose-viability curves as for NHEK cells but (depending on the cell model) varying susceptibility to oxidative products derived from RSV treatment (Figure 2c-d). Slight but significantly increased expression of molecular markers for oxidative stress response, such as catalase (CAT), could be observed up to 100 μM RSV with a maximum at 50 μM RSV (Figure 2b). On the other hand, too high concentrations of RSV (> 100 μM RSV) can result in toxic effects (Figure 2b). In summary, these data are in line with a large body of mostly unconsciously reported hormetic cellular effects of RSV ^10^.

**Figure 2.**
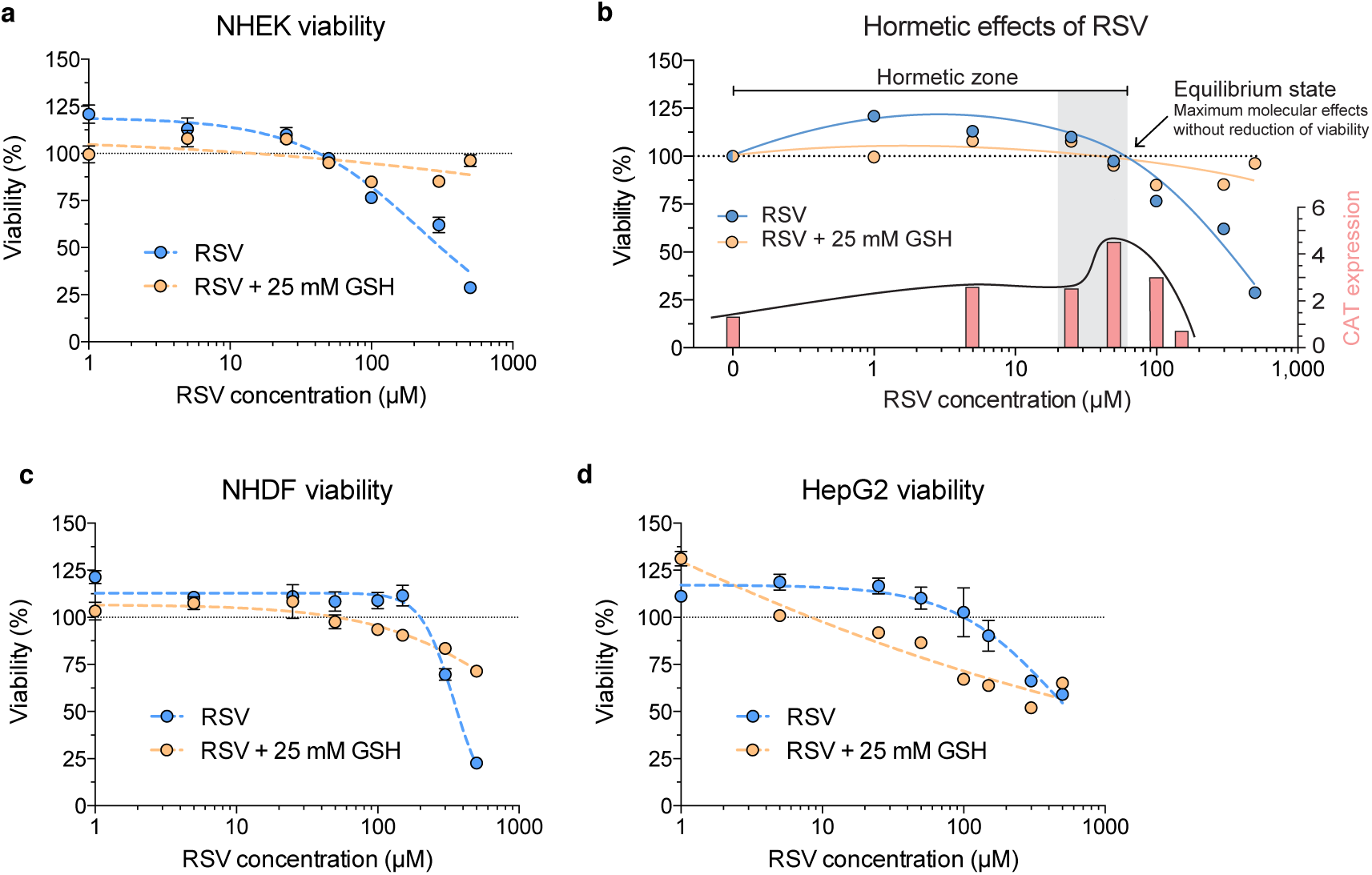
Hormetic effects are induced by generation of ROS derived from RSV. **(a)** RSV decreased viability of NHEKs after 16 hours treatment at concentrations ≥ 50 μM (IC_50_: 223.8 μM). RSV effects on viability are considerably quenched by GSH (IC_50_: 247,747.0 μM). Values are mean ± s.e.m. (n = 3). **(b)** Dose-response curve of RSV in treated NHEKs (blue). Hormetic zone emerges from 1 to 60 μM RSV and highest molecular effects (gene expression peaks of oxidative stress response gene catalase (CAT)) were generally observed at 50 μM RSV. **(c)** RSV decreased viability of NHDF cells at concentrations ≥ 300 μM (IC_50_: 342.5 μM), which was quenched by GSH (IC_50_: 1,163.0 μM). Values are mean ± s.e.m. (n = 6). **(d)** RSV decreased viability of human HepG2 liver cells at concentrations ≥ 150 μM (IC_50_: 445.3 μM), GSH quenched RSV induced effects. Values are mean ± s.e.m. (n = 6).

Importantly, the hormetic dose-viability curve was strongly truncated by adding 25 mM synthetic GSH as a quencher, providing to our knowledge for the first time strong evidence that increased viability of cells after RSV treatment mainly derived from ROS and related products of RSV (Figure 2a-d). In other words, the mechanism of action of RSV to slightly improve cellular fitness seems to rely significantly on oxidative effects of RSV, resulting in a bi-phasic, concentration-dependent cellular response.

Further concentration‐ and time-dependent treatments of NHEKs resulted in generally slight up-regulation of a number of metabolic, aging, oxidative stress and inflammation signalling genes (Figure 3a-d and Fig. S5e-h). Notably, in NHEKs these cellular responses could be observed in a concentration range of approximately 5 to 100 μM RSV. Strong changes in gene expression were observed after at least 12 hours, most efficiently after 16 hours of treatment.

**Figure 3.**
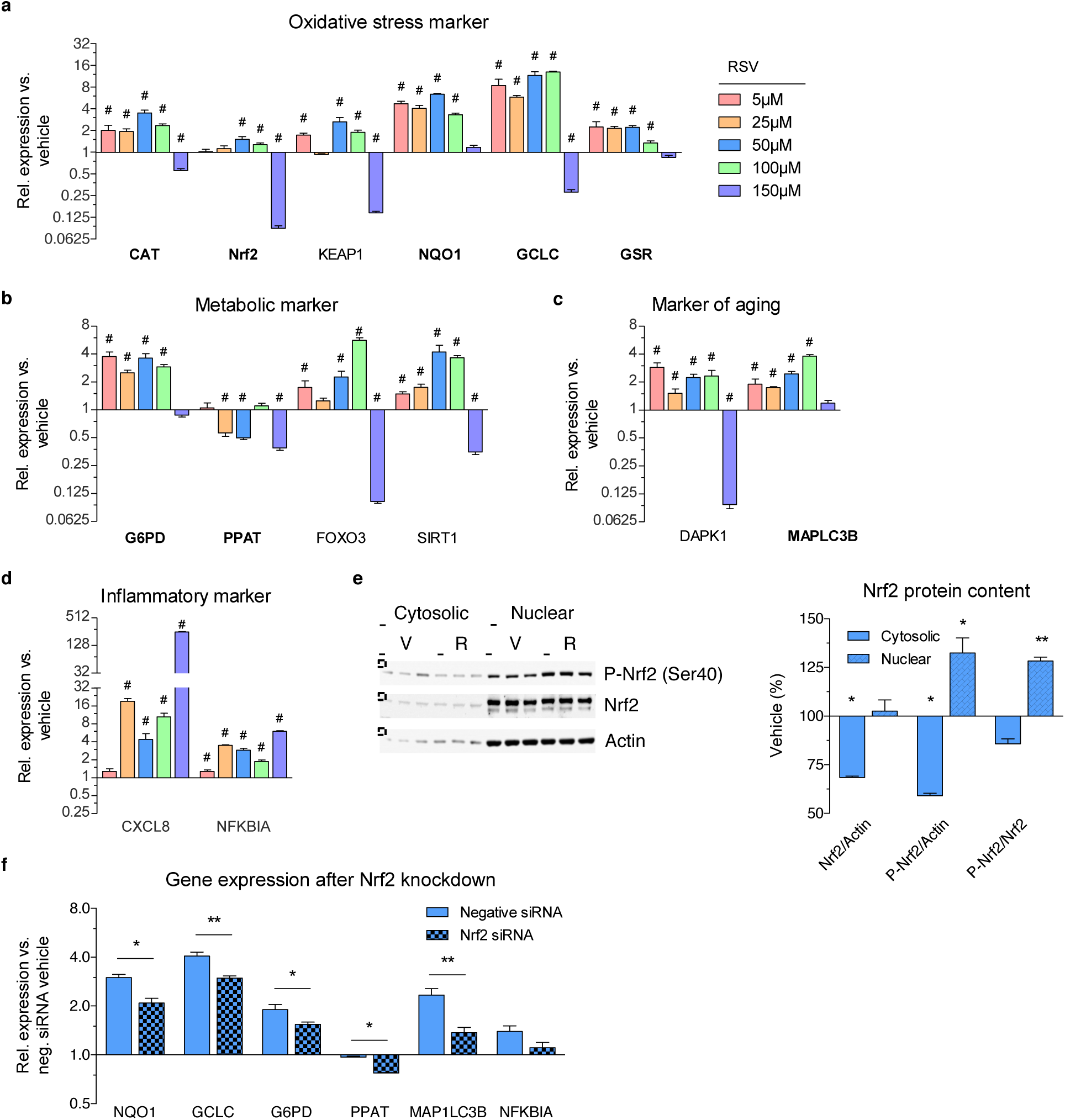
Cellular response to RSV-derived oxidation products including ROS is mediated by the redox-sensitive transcription factor Nrf2. **(a, b, c and d)** Concentration-dependent effects of RSV in NHEK cells after 16 hours. Values are mean ± s.e.m. (n = 4); # *P* < 0.05 versus vehicle. Nrf2 and its target genes are marked in bold font. See also Fig. S5a-d. **(e)** Translocation of Nrf2 and increased amount of nuclear phosphorylated Nrf2 as well as nuclear Nrf2 protein. Densitometric analysis of immunoblots relative to vehicle, with all gels run under the same experimental conditions and cropped blots depicted (copping: dashed line). Values are mean ± s.e.m. (n = 4), ** *P* < 0.01. V, vehicle; R, RSV.**(f)** Quantification of gene expression after knockdown of Nrf2 in RSV treated (16 hours) NHEKs. Values are mean ± s.e.m. (n = 4); * *P* < 0.05, ** *P* < 0.01, *** *P* ≤ 0.001 one-way ANOVA versus negative siRNA vehicle. See also Fig. S6b.

The entirety of the above shown data suggests that under physiological relevant conditions increased viability of cells after RSV treatment was triggered by ROS (and potentially also other radicals of RSV), leading to up-regulation of major cell defence genes. Instead of the conventional pharmacological (linear) threshold model we observed a bi-phasic mode of action of RSV: at normally applied non-toxic concentrations, RSV treatment results in increased cellular fitness based on related molecular events, whereas at higher concentrations RSV treatment results in toxic effects. The underlying reason for this cellular behaviour seems to depend largely on the pro-oxidative properties of RSV.

In summary, the here proposed link introduces an explicit explanation of the so far rather “nebulous” hormetic effects of RSV.

### Hormetic effects of oxidative products derived from RSV are driven by activation of Nrf2

We next asked how oxidising RSV could induce any specific molecular response in a cellular context. Especially the nuclear factor (erythroid-derived 2) like 2 (Nrf2) is considered responsible for accommodating oxidative stress ^2,29-31^. Consistent with the above shown production of oxidation products of RSV, we observed translocation of redox-sensitive Nrf2 into the nucleus of NHEKs (Figure 3e and Fig. S6a), leading to regulation of known Nrf2 target genes (Figure 5a-d bold font). Remarkably, knockdown of mRNA expression of Nrf2 gene by small interfering RNAs (siRNAs) significantly decreased the observed effects of RSV on gene expression response (Figure 3f and Fig. S6b). This experiment indicates that Nrf2 via its well-established canonical signalling model mediates major response of NHEKs to the oxidation products of RSV.

ROS and further radicals can produce numerous effects as a result of increased oxidation of cellular biomolecules, leading for example to inhibition of protein activity. Thus, we would expect multiple cellular defence mechanisms to counteract ROS including for example autophagy and cell cycle arrest. Interestingly, RSV treatment slightly increased autophagy, probably to degrade and recycle potentially damaged cellular components (Figure 4a and Fig. S6c), corroborating previous observations ^32^. Simultaneously, primary NHEKs were arrested in G1 cycle phase (Figure 4b and Fig. S6d). Interestingly, the cells did not show any signs of senescence or apoptosis (Figure 4c and d, Fig. S6e and f) and revealed reduced necrosis (Figure 4d).

**Figure 4.**
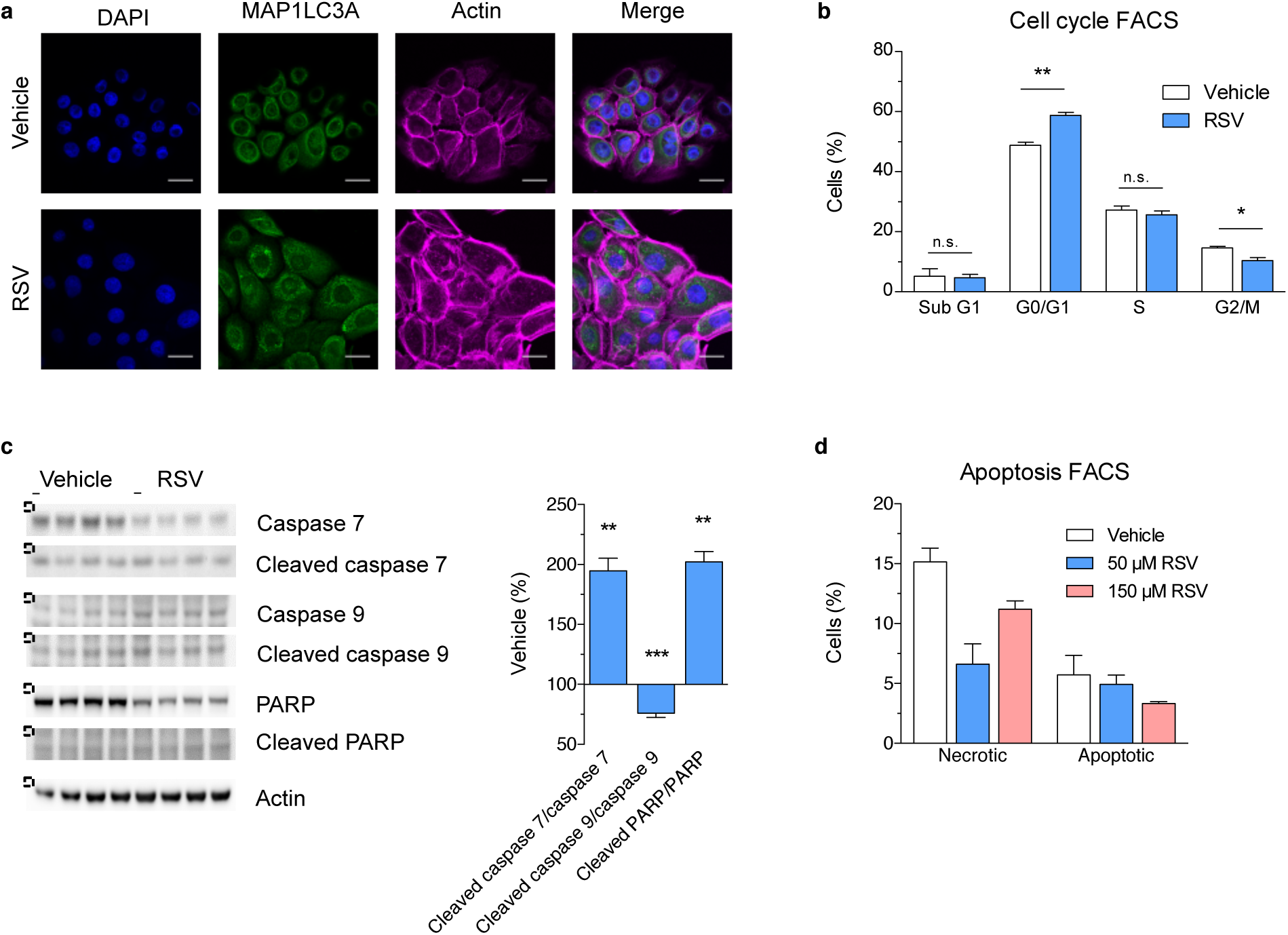
RSV induced autophagy and G1 phase cell cycle arrest but did not cause apoptosis in NHEK cells. **(a)** Fluorescence staining marking DAPI (blue), microtubule-associated protein 1 light chain 3A (MAP1LC3A, green) and Actin (magenta) in RSV treated NHEKs. See also Fig. S5c. **(b)** G1 phase cell cycle arrest in NHEKs after 16 hours of treatment with 50 μM RSV. Values are mean ± s.e.m. (n = 4); * *P* < 0.05; ** *P* < 0.01, *** *P* ≤ 0.001 versus vehicle. See also Fig. S6d. **(c)** RSV substantially increased the ratio of caspase 7 and PARP suggesting slight apoptosis. All gels were run under the same experimental conditions and cropped blots depicted (copping: dashed line). Values are mean ± s.e.m. (n = 4); * *P* < 0.05, ** *P* < 0.01, *** *P* ≤ 0.001 versus vehicle, **(d)** None of the tested RSV concentrations did induce apoptosis after 16 hours of treatment of NHEKs, but 150 μM RSV tend to increase necrotic cell proportion. Values are mean ± s.e.m. (n = 2). See also Fig. S6f.

In the context of increasing autophagy and overall molecular stress, this mechanism might allow cells to focus their limited resources on cellular repair, while decreasing cellular proliferation and nucleotide synthesis (Figure 1e). Notably, similar effects were observed for mild stress such as calorie restriction to improve cellular fitness ^33, 34^. Evidently, many of the effects described appear to be specifically mediated via activation of the redox-sensitive transcription factor Nrf2. We next asked if and how other reported factors such as the promiscuously reacting deacetylase SIRT1 could potentially modulate the effects derived from RSV-based activation of Nrf2, for example in the context of autophagy.

As shown above, under physiologically relevant conditions the almost completely degraded RSV can merely directly or allosterically induce the enzymatic activity of SIRT1 (Fig. S1e). However, the 4-fold up-regulation of SIRT1 expression (Figure 3b), the phosphorylated SIRT1 (Fig.S7a) and the simultaneously increased NAD^+^/NADH ratio (Figure 5a and Fig. S8d) might contribute to modify the mild effects derived from oxidized RSV. Knockdown of SIRT1 gene resulted in 20% lower expression of NRF2, suggesting a potential modifying effect of the lysine deacetylase SIRT1 on Nrf2 (Fig. S7b). However, at least in NHEKs – in contrast to knockdown of NRF2 – knockdown of SIRT1 did not strongly influence overall gene or protein expression (Fig. S7c and d), corroborating the major role of Nrf2 in the response to oxidative products such as ROS derived from RSV.

**Figure 5.**
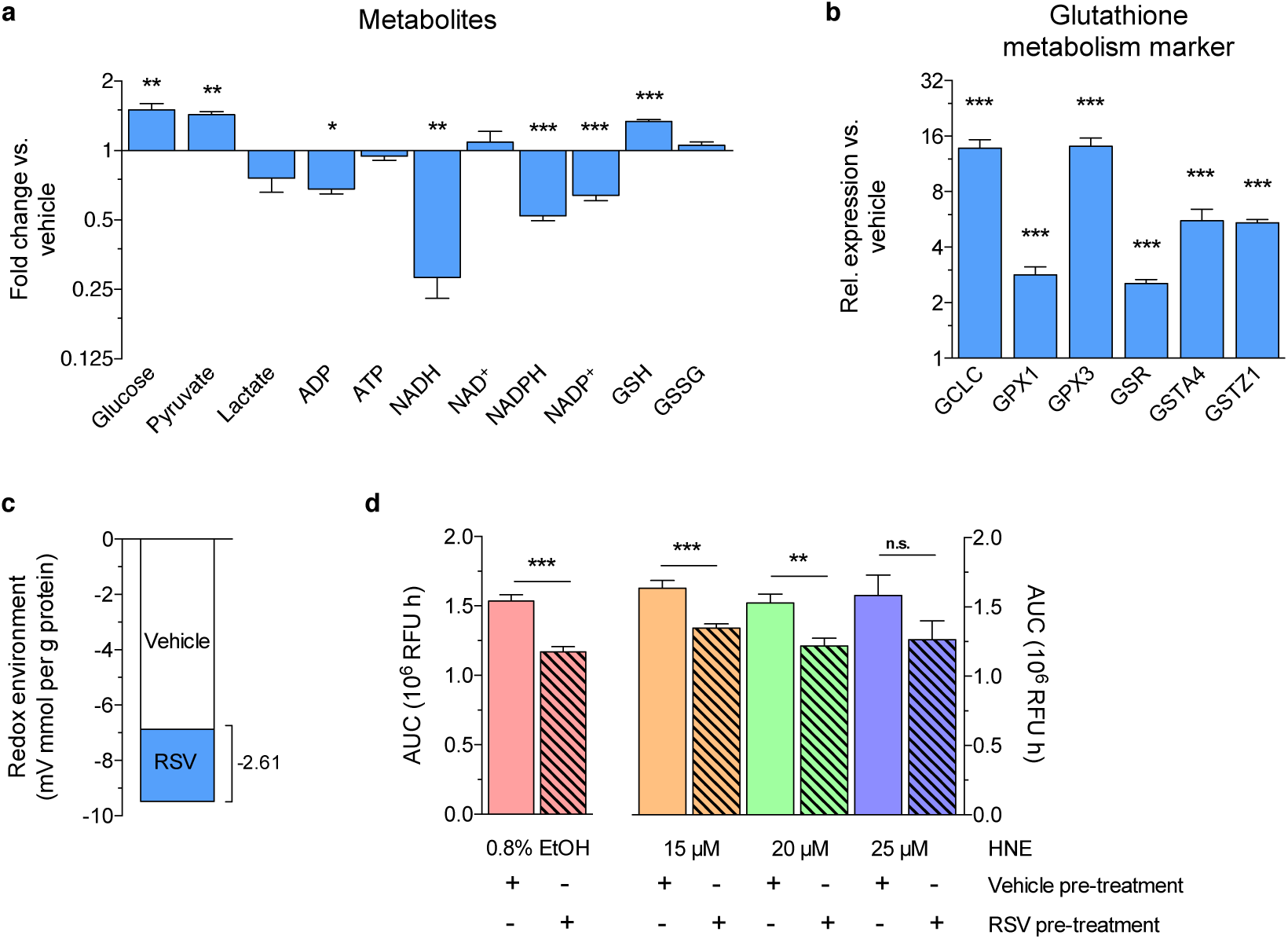
RSV influenced the concentration of key metabolites to shift the cellular redox environment to a reduced state, thereby increasing robustness versus oxidative stress. **(a)** Shift in intracellular metabolite concentrations after treatment with 50 μM RSV for 16 hours in NHEKs. Values are mean ± s.e.m. (NADH, NAD^+^ n = 3; glucose, ATP, ADP, GSH, GSSG n = 4; lactate, pyruvate, NADPH, NADP^+^ n = 5); * *P* < 0.05, ** *P* < 0.01; *** *P* ≤ 0.001 versus vehicle. See also Fig. S8d. (b) RSV substantially induced gene expression of glutathione metabolism marker genes. Values are mean ± s.e.m. (n = 4); *** *P* ≤ 0.001 versus vehicle. **(c)** 2GSH/GSSG ratio was used to calculate the redox environment. See also Table S4 and for calculation with various metabolites Fig. S8f. (d) Pretreatment of NHEKs with 50 <M RSV for 16 hours decreased intracellular ROS generation after subsequent treatment with ethanol (0.78 %), while increasing concentrations of thiol/GSH-scavenger HNE revised ROS-protective potential. Values are mean ± s.e.m. (n = 7); ** *P* < 0.01, *** *P* ≤ 0.001 versus vehicle. AUC, area under the curve; RFU, relative fluorescence units.

### Oxidative products of RSV induce a reduced cellular redox environment

We then asked how gene expression mediated by RSV/ROS-based activation of Nrf2 might influence cellular metabolism. In NHEKs treated for 16 hours with 50 μM RSV, we observed phosphorylation signalling events such as increased phosphorylation of pyruvate dehydrogenase E1 component subunit alpha (PDE1α) at serine 293 (Fig. S8a), an effect known to inhibit endogenous pyruvate breakdown and oxidative phosphorylation ^35^. Consequently, this molecular effect led to increased intracellular levels of the potential ROS-scavenger pyruvate while lactate remained at constant levels (Figure 5a). Under constant mitochondrial biogenesis (Fig. S8b), we further observed decreased mitochondrial oxygen consumption during 16 hours of RSV treatment (Fig. S8c).

Moreover, the intracellular ratio of metabolite couples 2GSH/GSSG and ATP/ADP as well as the amount of glucose significantly increased (Figure 5a and Fig. S8d). In contrast the intracellular ratios of metabolite couples NADH/NAD^+^ and NADPH/NADP^+^ were significantly decreased (Figure 5a and Fig. S8d). Consistent with the ratio of the most relevant 2GSH/GSSG redox couple, the expression of genes and proteins related to glutathione metabolism were highly increased, corresponding to significantly elevated levels of the potent cellular antioxidant GSH (Figure 5a and b, Fig. S8d and e).

In summary, these data indicate a metabolic switch that leads amongst others to an increased pool of reduced glutathione. Clearly, the GSH concentration can vary a lot between different cells, depending on stress exposure and function ^36^.

We next analysed the redox environment of NHEKs treated with RSV using intracellular concentrations of above-mentioned key metabolites, using the formula (Eq. 1) ^36^:

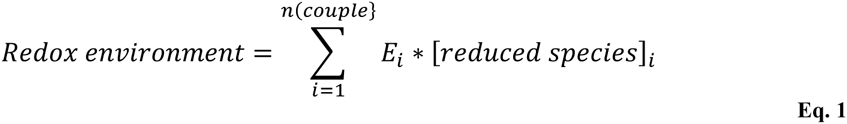

Indeed, RSV treatment at hormetic concentrations shifted the cellular redox environment to a more reduced state mediated by Nrf2 (Figure 5c and for calculation-relevant parameters Table S4) ^37^. Based on the 2GSH/GSSG couple, which provides the by far largest pool of reducing equivalents ^36^, we calculated a slight shift of redox environment of -2.61 mV mmol GSH per gram protein. A similar trend was observed by taking into account further redox couples (see also Fig. S8f and Table S4). According to our results, RSV treatment can contribute to an overall reduction of biological molecules containing for example thiol groups, as evident from the increased cellular GSH concentration (Figure 5a).

We then analysed if the observed reduced redox environment could potentially protect the cell from (oxidative) stress. Therefore, we subjected NHEKs to 16 hours pre-treatment with 50 μM RSV. After replacement of medium the NHEK cells were devoid of any residual RSV. We then treated NHEKs with ~0.8% ethanol and analysed the endogenous generation of intracellular ROS from cellular metabolisation of ethanol, i.e. the level of oxidative stress (Figure 5d)^38^. Notably, we observed that overall reduced cellular environment (owing to the observed increased pool of endogenous GSH) enabled RSV-pre-treated NHEKs to buffer the additional production of ROS due to biotransformation of ethanol (Figure 5d). The protective effects of pre-treatment with RSV were revised in a concentration-dependent manner by addition of 4-hydroxy-2-nonenal (HNE), an α, β-unsaturated aldehyde, which amongst others acts as a strong electrophile by depleting cellular sulfhydryl compounds like GSH (Figure 5d) ^39^.

Summarizing, we propose a bi-phasic pharmacological model for RSV, which might be extended to other pro-oxidative polyphenols. This model comprises i) generation of oxidation products at non-toxic concentrations in physiologically relevant sodium bicarbonate-containing media, ii) specific mediation of cellular response to oxidation induced by RSV by the redox-sensitive transcription factor Nrf2 and iii) induction of slight reductive shifting of cellular redox-environment to protect the cell from additional (oxidative) stress (Figure 6a and b).

**Figure 6.**
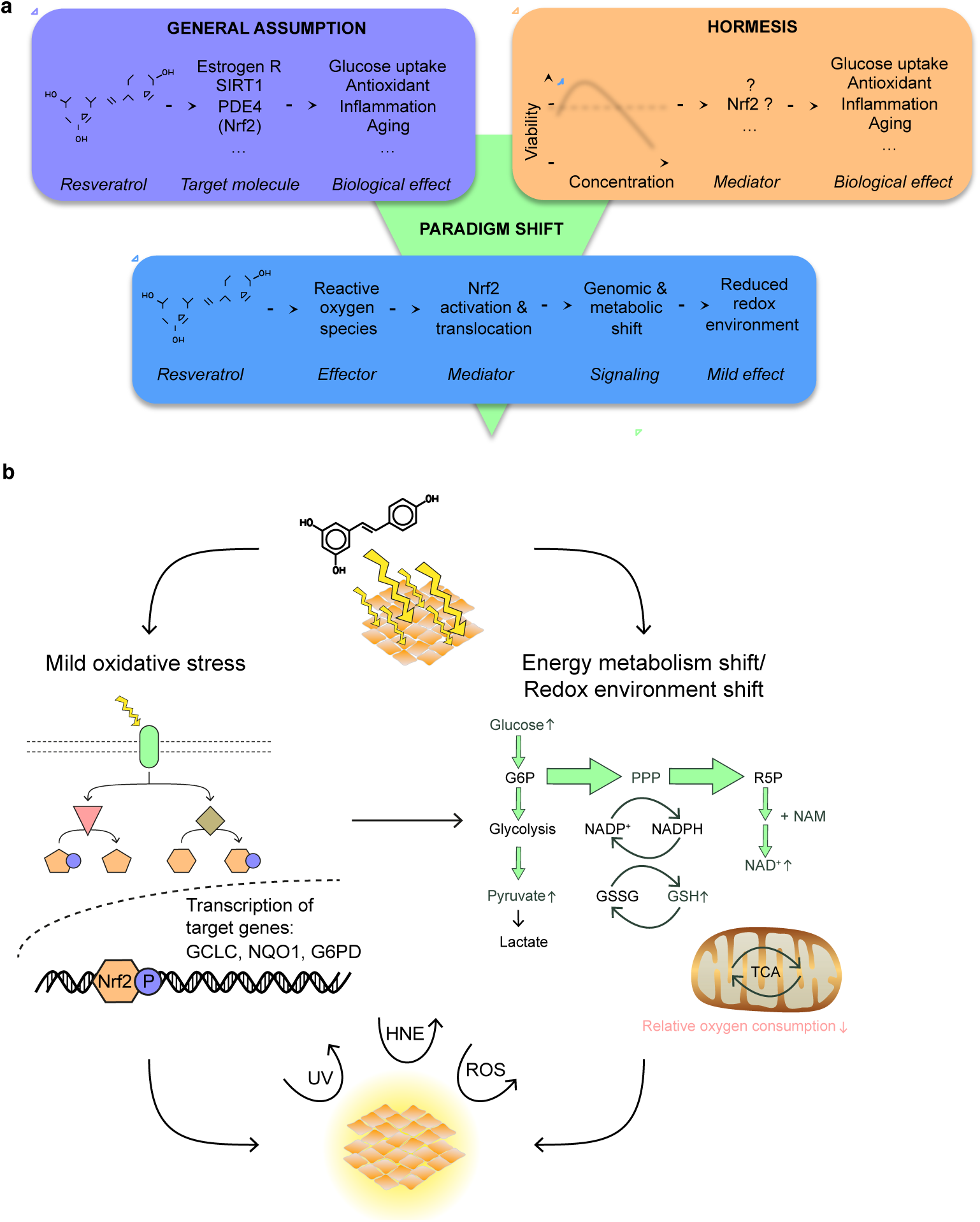
Molecular networks involved in RSV-induced cytoprotection against oxidative stress, **(a)** A dose-dependent compound-target protein model is generally assumed to explain biological effects of RSV. Our proposed model integrates mechanism of action of RSV via production of ROS in physiologically relevant media, Nrf2-mediated cellular response and mild beneficial (hormetic) biological effects at normally applied concentrations. Estrogen R = estrogen receptor, SIRT1 = silent mating type information regulation 2 homolog 1, PDE4 = phosphodiesterase 4. **(b)** At low concentrations oxidising RSV induces mild stress, resulting in a metabolic and redox environment shift to increase the level of endogenous quenchers to eventually protect the cell from environmental stress.

Clearly, the here proposed model largely depends on the pro‐ and anti-oxidative properties of RSV in a given microenvironment. We argue that bicarbonate containing media are essential for living systems, and thus it seems that our model might be extendable to a large variety of biological phenomena. Our data indicate at least that proper analysis of the potential pro‐ and anti-oxidative features of RSV in a specific experimental set-up shall be thoroughly explored prior any biological investigation to define a common basis and thereby avoid any potential misinterpretation.

In particular the human epidermis being predominantly prone to external stress might benefit from dermatological application of RSV. The here elaborated pro-oxidative features of RSV and the redox-environment shifting concept can fully explain such recent claims ^31^. Of note, whereas RSV might remain stable under low pH conditions in the stomach (Fig. S2 and 3), physiologically well-known neutralization in the duodenum – by high concentrations of bicarbonate derived from the pancreas – might induce so far largely unexplored pro-oxidative features of RSV in the intestine.

Considering reduction of the cellular redox environment as the main physicochemical mechanism, the often-reported weak and pleiotropic effects of RSV can be quantitatively determined using molecular data of redox-relevant metabolites and the above formula (Eq. 1; see also Figure 5c and Table S4). On the other hand, in particular the effects derived from oxidation products with short lifetimes such as ROS are difficult to trace *in vitro* and in particular *in vivo*. This drawback might provoke scepticism how such “dirty” chemicals shall exert well-controllable cellular and physiological effects.

Nevertheless, taking a systems view we suggest to apply the here explored paradigm for mathematically analysing the emerging relevant biological effects of “dirty” compounds. This approach allows amongst others calculating the oxidation-buffering capacity of targeted cells under investigation. Using such a framework, development of chemical derivatives of RSV and of molecular carriers might be helpful to rationally exploit the beneficial bi-phasic effects of pro-oxidative compounds for therapeutic and preventive application.

In general, hormetic induction of cellular fitness by physiologically pro-oxidative polyphenols such as RSV might represent a powerful approach to protect cells against physiological stress and to inhibit age-related diseases.

## Methods

### Materials

Chemical compounds were purchased from the following sources: 3,5,4'-trihydroxy-*trans*-stilbene (resveratrol, RSV) and 4-hydroxy-2-nonenal (HNE) from Cayman Chemical (Biomol, Hamburg, Germany). Reduced glutathione (GSH) was purchased from Sigma Aldrich (Taufkirchen, Germany). The composition of Berlin tap water can be retrieved from: http://www.bwb.de

### Cell culture

*Neonatal normal human epidermal keratinocyte cells* (NHEK, CC-2503, Lonza, Basel, Swiss) were isolated from a black, newborn male. NHEK cells were maintained in keratinocyte growth medium (KGM) containing keratinocyte basal medium (KGM, CC-3101, Lonza) and KGM SingleQuot Kit Suppl. & Growth Factors (CC-4131, Lonza). Cells were treated with indicated compounds, vehicle controls were as follows: DMSO for RSV and EtOH for HNE. GSH was dissolved in cell culture medium. Notably, hydrogen peroxide applied in low micromolar concentrations (comparable to HNE) did not produce any significant oxidative effects, probably due to efficient scavenging in the cellular context. *Neonatal normal human dermal fibroblast cells* (NHDF, CC-209, Lonza) were isolated from a Caucasian, newborn male. NHDF cells were maintained in Dulbecco’s modified Eagle medium (# 31966, Gibco, Life Technologies, Darmstadt, Germany) supplemented with 10% fetal bovine serum (FBS) Medium was renewed every two to three days and cells were split two times per week. Cells were treated at 60% confluence with 50 μM RSV or vehicle control. *Human HT-29 colon cells* (ACC-299, DSMZ, Braunschweig, Germany) were cultured in Dulbecco's Modified Eagle Medium/Nutrient Mixture F-12(DMEM/F-12, # 11330-057, Gibco, Life Technologies) supplemented with 5% and 100 U/ml penicillin and 100 μg/ml streptomycin (all Biochrom, Berlin, Germany). *Human THP-1 monocyte cells* (ATCC, LGC Standards GmbH, Wesel, Germany) were cultivated in RPMI 1640 (Biochrom) supplemented with 10% FBS and 100 U/ml penicillin and 100 μg/ml streptomycin. *Human HepG2 liver cells* (ATCC, LGC Standards GmbH), human embryonic kidney (HEK293T) cells (ATCC, LGC Standards GmbH) and *human HeLa cells* (ATCC, LGC Standards GmbH) were cultured in DMEM (# 31966, Gibco, Life Technologies) supplemented with 10% FBS and 100 U/ml penicillin and 100 μg/ml streptomycin. Cells were seeded into 12-well plates (# 3513, Corning, Fisher Scientific, Schwerte, Germany) and treated with RSV or vehicle control. Human adult low calcium high temperature keratinocyte cells (HaCaT) and ARE clone 7 HaCaT cells were kindly provided by Unilever (Sharnbrook, U.K.). ARE clone 7 HaCaT cells were revived in DMEM (# 31966, Gibco, Life Technologies) supplemented with 10% FBS. The next day, culture medium was changed to selective medium (DMEM with 10% FBS and 400 μg/ml Hygromicin B (# 10687-010, Life Technologies)) for continued growth. HaCaT cells were cultured in DMEM (# 21068-028, Life Technologies) supplemented with 1% FBS, 2 mM L-glutamine (Biochrom), 1 mM sodium pyruvate (# 11360-039, Life Technologies), 70 μM calcium chloride (Merck GmbH, Darmstadt, Germany), 100 U/ml penicillin and 100 μg/ml streptomycin.

All cell lines were maintained at 37°C in a humidified 5% CO_2_ atmosphere and treated at 60% confluence. The following passages were used: NHEK cells p 1, NHDF cells p 2-5, ARE clone 7 HaCaT cells p 7, HaCaT cells p 47, HT-29 cells p 27, HepG2 cells p 9, HEK293T p 23 and Hela were kindly provided by Dr. David Meierhofer.

### Analysis of resveratrol integrity

The Fluor-de-Lys SIRT1 Fluorometric Drug Discovery Assay (BML-AK555-0001, Enzo Life Sciences, Lörrach, Germany) was used to analyse the integrity of RSV after incubation in diverse solvents according to manufacturer’s instructions. Briefly, compounds were added to a mixture of NAD^+^ and Fluor-de-Lys (FdL) peptide in reaction buffer (RB). Reaction was started by addition of SIRT1 in RB followed by 30 min incubation at 37°C. Reaction was stopped by addition of NAM and DeveloperII solution. After additional incubation for 45 min at 37°C, fluorescence was measured (360/460 nm) using the POLARstar Omega (BMG LABTECH, Ortenberg, Germany).

### Time-dependent decay of resveratrol (cell-free)

The time-dependent decay of RSV in diverse solvents was analysed using the POLARstar Omega (BMG LABTECH) at 37°C. Samples were transferred (100 μl/well) into a UV-Star 96-well plate (# 655801, Greiner Bio-one, Frickenhausen, Germany) for kinetic and spectral measurement (between 220 and 720 nm, Δλ 2 nm). Photos of RSV under various conditions (Fig. S1a and b) to document initial decay overnight were taken after about 17.5 hours.

### pH-dependent oxidation of resveratrol (cell-free)

The time-dependent oxidation of 50 μM RSV in ddH_2_O with or without 44 mM sodium bicarbonate (NaHCO_3_) was analysed using the POLARstar Omega (BMG LABTECH) at 37°C. Samples were transferred (150 μl/well) into a UV-Star 96-well plate (# 655801, Greiner Bio-one) for kinetic and spectral measurement (between 230 and 550 nm, Δλ 2 nm). The pH of each solution was adjusted from 1 to 12 using HCl and NaOH solutions. In accordance to Li, et al. ^14^ oxidation products of RSV, a shortlived hydroxyl radical adduct of RSV (characteristic absorbance maximum: 420 nm) and the relatively stable 4’-phenoxyoxyl radical (characteristic absorbance maximum: 390 nm), were monitored. For data analyses in GraphPad Prism 5.0 signals were background-subtracted and normalized to vehicle control. Data were fitted (dashed line) using GraphPad Prism 5.0 with a Hill slope of -1 according to equation:

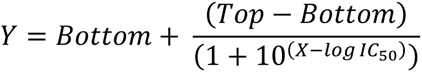

### Oxygen partial pressure-dependent oxidation of resveratrol (cell-free)

96-well plates prepared for the determination of the pH-dependent oxidation of resveratrol (see pH-dependent oxidation of RSV) were incubated at 37°C at atmospheric oxygen levels (~21% O_2_), slightly reduced oxygen partial pressure (10% O_2_, mimicking conditions in the blood vessels), or highly reduced oxygen levels (1% O_2_, resembling tissue or tumour microenvironment). For experiments with reduced oxygen partial pressure, plates were incubated at corresponding oxygen levels using a CO_2_ Incubator Model CB 60 (Binder, Tuttlingen, Germany). For spectral measurements plates were quickly analysed (< 2 min) using the POLARstar Omega (BMG LABTECH) at 37°C. Afterwards the plates were further incubated at indicated conditions. In accordance to Li, et al. ^14^ the oxidation of RSV and subsequent reaction products were monitored. For data analyses in GraphPad Prism 5.0 signals were background-subtracted and normalized to vehicle control. Data were fitted (dashed line) using GraphPad Prism 5.0 with Hill slope = -1 according to equation:

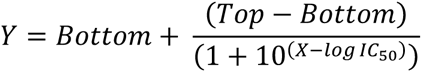

### Measurement of ROS (cell-free)

The CellROX Green dye (C10444, Life Technologies) was used to quantify the formation of extracellular ROS. The dye exhibits a high fluorescence response in particular to hydroxyl radicals (OH^-^). Upon oxidation by ROS and after binding to nucleic acid the probe exhibits green photostable fluorescence. The CellROX Green dye is compatible with cell culture medium and requires no cellular processing, hence it is applicable for measurement of ROS generation in a cell-free environment. The dye was diluted to 10 μM/well in presence of 1 μg/ml lambda DNA (Life Technologies) prior to addition of the indicated compounds. Measurement was performed in a final volume of 150 μl/well in black 96-well plates (# 655090, Greiner Bio-One). The dye was protected against atmospheric oxygen by adding a sealing layer of 100 μl/well mineral oil (Luxcel Biosciences). Fluorescence intensity (Ex 485/30; Em 530/10) was recorded at 37°C for 16 hours using the POLARstar Omega (BMG LABTECH). For data analyses in GraphPad Prism 5.0 fluorescence signals were background-subtracted and normalized to vehicle control. Finally, signals were plotted in GraphPad Prism 5.0 either using a second order neighbour smoothing (4 neighbours) for kinetic depiction or the area under the curve (AUC) was calculated for summed depiction.

### Quantification of intracellular ROS

Formation of ROS was quantified using dye 5-(and-6)-chloromethyl-2',7'‐ dichlorodihydrofluorescein diacetate (CM-H2DCFDA, Life Technologies) according to the manufacturer’s instruction. In detail, NHEK cells were seeded in a 96-well plate (TPP, Biochrom) with a density of 10,000 cells/well. The following day, adherent cells were washed once with pre-warmed PBS and loaded with 50 μM dye diluted in PBS. For successful incorporation and activation of CM-H_2_DCFDA, cells were incubated for 30 minutes at 37°C. Afterwards, free dye was removed by washing with pre-warmed PBS. KGM (100 μl/well) was added and cells were once more incubated at 37°C for 60 minutes. Compounds were added as indicated and fluorescence (Ex 485/30; Em 530/10) was measured for 16 hours of treatment using the POLARstar Omega (BMG LABTECH) at 37°C.

For the quantification of ROS generation in pre-conditioned NHEKs, cells were seeded in a 96-well plate (TPP, Biochrom) with a density of 10,000 cells/well and were pre-treated with 50 μM RSV or vehicle for 16 hours. The following day, adherent cells were washed once with pre-warmed PBS and loaded with 50 μM dye diluted in PBS. For successful incorporation and activation of CM-H_2_DCFDA, cells were incubated for 30 min at 37°C. Afterwards, free dye was removed by washing with pre-warmed PBS. KGM (100 μl/well) was added and cells were once more incubated at 37°C for 60 min. Putative protection of NHEKs against oxidative stress by RSV pre-treatment was tested by adding ethanol (0.781%) or potent thiol-scavenger HNE ^40^ at indicated concentrations. Fluorescence (Ex 485/30; Em 530/10) was measured for 21 hours of treatment using the POLARstar Omega (BMG LABTECH) at 37°C. For data analyses in GraphPad Prism 5.0 fluorescence signals were background-subtracted and normalized to vehicle control. Finally, signals were plotted in GraphPad Prism 5.0 either using a second order neighbour smoothing (4 neighbours) for kinetic for kinetic depiction or the area under the curve (AUC) was calculated for summed depiction.

### Analysis of H_2_O_2_ generation (cell-free)

To determine hydrogen peroxide (H_2_O_2_) generation the Amplex Red Glucose/Glucose Oxidase Assay Kit (A22189, Life Technologies) and the Hydrogen Peroxide Assay Kit (K265-200, BioVision, BioCat, Heidelberg, Germany) were used. DMEM samples were filtered using Amicon Ultra-0.5 Centrifugal Filter Unit with Ultracel-10 membrane (UFC501024) or MultiScreen Ultracel-10 Filter Plate 10 kD (MAUF01010, both Merck Chemicals). For each experiment a H_2_O_2_ standard curve was generated. Samples were mixed with provided dye, horseradish peroxidase (HRP) and RB. Fluorescence was measured after 10 to 30 minutes incubation at RT using the POLARstar Omega (BMG LABTECH).

### Analysis of superoxide generation (cell-free)

The MitoSOX Red Mitochondrial Superoxide Indicator (M36008, Life Technologies) was used to quantify the formation of superoxide. The dye is readily and specifically oxidized by superoxide and exhibits red photostable fluorescence after binding to nucleic acids. The probe is compatible with cell culture medium and in combination with Lamda DNA (Life Technologies) applicable for measurement of superoxide generation in a cell-free environment. The dye was diluted to 10 μM/well in presence of 200 ng/well lambda DNA (Life Technologies) prior to addition of indicated compounds. Measurement was performed in a final volume of 150 μl/well in black 96-well plates (# 655090, Greiner Bio-One). The dye was protected against atmospheric oxygen by adding a sealing layer of 100 μl/well mineral oil (Luxcel Biosciences). Fluorescence intensity (Ex 485/30; Em 530/10) was recorded at 37°C for 16 hours using the POLARstar Omega (BMG LABTECH). For data analyses in GraphPad Prism 5.0 fluorescence signals were background-subtracted and normalized to vehicle control. Finally, signals were plotted in GraphPad Prism 5.0 either using a second order neighbour smoothing (4 neighbours).

### Viability assay

For determination of cellular viability NHEK cells were seeded in a black 96-well plate (# 353219, BD Biosciences, Heidelberg, Germany) with a density of 10,000 cells/well and a final volume of 200 μl/well. NHDF, HepG2 and THP-1 cells were seeded in a black 384-well plate (# 3712, Corning, Fisher Scientific) with a density of 2,500 cells/well (NHDF) and 5,000 cells/well (HepG2, THP-1), respectively and a final volume of 25 μl/well. The following day, medium was renewed and cells were treated with the indicated compounds in a final volume of 100 μl/well (NHEK) and 35 μl/well (NHDF, HepG2, THP-1), respectively. After 16 hours of treatment, cell viability was quantified using the CellTiter-Fluor Cell Viability Assay (G6081, Promega, Mannheim, Germany). Fluorescence intensity was measured (410/520 nm) using the POLARstar Omega (BMG LABTECH). Data were fitted (dashed line) using GraphPad Prism 5.0 with variable Hill slope according to equation:

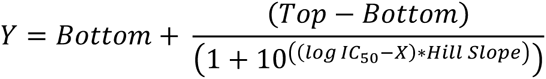

Fluorescence intensity values were transformed to the relative number of cells. IC_50_ is the concentration required for a 50 % inhibition of viability

### RNA isolation, reverse transcription and quantitative real-time PCR

RNeasy Mini Kit (QIAGEN, Hilden, Germany) was used to isolate total RNA according to the manufacturer’s instruction. For cell lysis 10 μl/ml ß-mercaptoethanolwas added to RLT buffer. Genomic DNA was digested on a column using the DNase-Set (QIAGEN). The concentration of extracted RNA was measured using the Nanodrop ND-2000 Spectrophotometer (Thermo Fisher Scientific). RNA was reversely transcribed into cDNA applying the High Capacity cDNA Reverse Transcription Kit (Life Technologies) with random primers. After an initial denaturation at 95°C for 10 min, the cDNA was amplified by 40 cycles of PCR (95°C, 15 sec; 60°C, 60 sec). Quantitative PCR was carried out on the ABI Prism 7900HT Sequence Detection System using the Power SYBR Green PCR Master Mix (all Life Technologies). The relative gene expression levels were normalized using ß-actin gene and quantified by the 2^-ΔΔCt^ method ^41^. Primer sequences are summarized in Table S5. Data were analysed using GraphPad Prism 5.0.

### Knockdown of NRF2 or SIRT1 with small interfering RNA

NHEK cells were seeded into 6-well plates (Corning) and transfected with 30 nM Silencer Pre-designed siRNA Nrf2 (# 16708), Silencer Pre-designed siRNA SIRT1 (# 136457) or Silencer Select negative control siRNA (# 4390844, all Ambion, Life Technologies) using Lipofectamin 2000 transfection reagent (# 11668019, Life Technologies). Transfection was carried out in 1 ml for 48 hours in KGM, whereby 0.5 ml KGM were added after 24 hours. Medium was then renewed and cells were treated with 50 μM RSV or vehicle for 16 hours prior to RNA and protein collection. Data were analysed using GraphPad Prism 5.0.

### Genome-wide gene expression analyses

Genome-wide gene expression analyses were done by ATLAS Biolabs GmbH (Berlin, Germany) on HumanHT-12 Expression BeadChips (Illumina, Eindhoven, The Netherlands). All basic expression data analyses were carried out using GenomeStudio V2011.1 (Illumina). Raw data were background-subtracted and normalized using the cubic spline algorithm. Processed data were filtered for significant detection (*P* value ≤ 0.01) and differential expression vs. vehicle treatment according to the Illumina t-test error model and were corrected according to the Benjamini-Hochberg procedure (*P* value ≤ 0.05) in the GenomeStudio software. Gene expression data were submitted to the Gene Expression Omnibus database (GSE72119).

Gene Set Enrichment Analysis (GSEA)^42^ of the RSV gene expression profiles was performed using the following parameters: 1000 gene set permutations, weighted enrichment statistic, and signal-to-noise metric. Microarray data were analysed using the curated C2 KEGG pathways gene sets (version 4.0, 186 gene sets) from the Molecular Signature Database (MSigDB) (Table S2).

### Cell cycle analyses

Analyses of cell cycle regulation were performed in NHEK cells treated with the indicated compounds for 16 hours. Trypsinised cells were fixed in 70 % ethanol and incubated on ice for 15 minutes. Fixed cells were then resuspendend in propidium iodide (PI)/RNase staining solution (# 4087, Cell Signaling Technology, New England Biolabs, Frankfurt, Germany), incubated for 15 minutes at RT and stored at -20°C until use. Finally, cells were measured in the Accuri C6 flow cytometer (BD Biosciences). Data analyses were performed using the Watson pragmatic model in FlowJo 7.6 (Tree Star).

### Phosphatidylserine externalization (Annexin V) assay

To quantify apoptosis, the externalization of phosphatidylserine ^5^ was determined in NHEK cells treated with the indicated compounds for 16 hours by staining with annexin-V-FLUOS and propidium iodide using the Annexin-V-FLUOS Staining Kit (Roche Diagnostics, Mannheim, Germany) and subsequent flow cytometry (Accuri C6, BD Biosciences). Analyses were performed using FlowJo 7.6 (Tree Star).

### Immunoblotting

NHEK were lysed in 50 mM Tris-HCl (pH 8.0), 10 mM EDTA, 1% SDS with protease inhibitors (Roche Diagnostics) and phosphatase inhibitors (Sigma Aldrich), and sonicated (Bandelin electronic, Berlin, Germany). After centrifugation for 10 minutes at 10,000 g, the supernatants were stored at -80°C until use. Samples were denaturated and separated using a NuPAGE Novex 4-12% Bis-Tris gel (Life Technologies) and blotted onto Hybond ECL nitrocellulose membranes (GE Healthcare, Freiburg, Germany). Membranes were blocked for 1 hour at RT according to the manufacturer's protocol and washed in TBS-T (0.1 %). Primary antibodies against P-Nrf2 (ab76026, Abcam, Cambridge, UK), Nrf2 (sc-13032X, Santa Cruz, Heidelberg, Germany), Sirt1 (MAb-063-050, Diagenode, Seraing (Ougrée), Belgium), P-Sirt1 (# 2314), CDK 4 (# 2906), cyclin D1 (# 2926), CDK 2 (# 2546), cyclin E2 (# 4132), p21 (# 2947), cyclin A2 (# 4656), caspase 7 (# 9492), cleaved caspase 7 (# 8438), caspase 9 (# 9502), cleaved caspase 9 (# 7237), PARP (# 9542), cleaved PARP (# 5625, all from Cell Signaling Technology) and b-Actin (sc-47778, Santa Cruz) were diluted in TBS-T (0.1%) with milk powder and BSA, respectively, according to the manufacturer's protocols. Membranes were shaken at 4°C overnight, washed 3 times with TBS-T (0.1 %) and subsequently incubated with anti-rabbit IgG-HRP (sc-2004, Santa Cruz) and anti-mouse IgG-HRP (sc-2005, Santa Cruz), respectively, according to the manufacturer's protocols. Detection was carried out with Western Lightning ECL solution (Perkin Elmer, Rodgau, Germany). Membranes were stripped with Restore Plus Western Blot Stripping Buffer (Thermo Scientific, Life Technologies) for 5 to 7 minutes. A densitometric analysis was performed with FUSION-SL Advance 4.2 MP System (Peqlab, Erlangen, Germany).

### Detection of lipid peroxidation

Click-iT Lipid Peroxidation Imaging Kit-Alexa Fluor 488 (C10446, Life Technologies) was used to determine the development of cellular lipid peroxides. Linoleamide alkyne (LAA, alkyne-modified linoleic acid) is incorporated into cellular membranes and oxidized upon lipid peroxidation and finally, resulting in proteins labelled with an azide-modified Alexa Fluor 488 dye. Increasing fluorescence intensities corresponds to enhance lipid peroxidation upon treatment. NHEK cells were seeded in 25 cm^2^ cell culture flasks (Corning) and treated with 50 μM RSV or vehicle for 16 hours in presence of 50 μM LAA. After trypsinisation, cells were washed with PBS and fixed in 3.7 % formaldehyde for 15 minutes at RT. Cells were washed in PBS, permeabilised by use of 0.5 % Triton X-100 in PBS for 10 minutes and subsequently blocked by adding 1 % BSA for 30 minutes (all at RT). Remaining BSA was removed by rigorously washing the cells with PBS. Pelleted cells were then incubated with 500 μL Click-iT reaction cocktail for 30 minutes at RT according to the manufacturer’s protocol. The Click reaction was stopped by adding 1 % BSA/PBS. The cells were washed and resuspended with PBS. Flow cytometry was performed on the Accuri C6 (BD Biosciences). Data were analysed using FlowJo 7.6 (Tree Star) and GraphPad Prism 5.0.

### Antioxidant Assay

The Antioxidant Assay (# 709001, Cayman Chemical) was conducted according to the manufacturer’s manual. The assay was miniaturized to a final volume of 60 μl/well. Samples including 6-hydroxy-2,5,7,8-tetramethylchroman-2-carboxylic acid (Trolox) standards were mixed with metmyoglobin and chromogen. The reaction was initiated by adding hydrogen peroxide solution and incubated 5 minutes at RT on a shaker. Finally, the absorbance was read at 405 nm with the POLARstar Omega (BMG LABTECH). Data were analysed using GraphPad Prism 5.0.

### Quenching of resveratrol effects

NHEK cells were seeded in 150 cm^2^ cell culture flasks (Corning) and treated with 50 μM RSV, vehicle alone or in combination with indicated concentrations of GSH, respectively for 16 h. Trypsinised cells were subjected to quantitative real-time PCR and 25 mM GSH measured in the Accuri C6 flow cytometer (BD Biosciences). Data were analysed using GraphPad Prism 5.0 and FlowJo 7.6 (Tree Star).

### Intracellular glucose quantification

Intracellular glucose concentration was determined using the PicoProbe Glucose Fluorometric Assay Kit (K688-100, Biovision, BioCat). NHEK cells were seeded into 25 cm^2^ cell culture flasks (Corning, Fisher Scientific) and treated with 50 μM RSV or vehicle for 16 h. Samples were processed according to the manufacturer’s instruction and deproteinised using the Deproteinising Sample Preparation Kit (K808-200, Biovision, BioCat). The assay was miniaturized to 10% of the initial volume and conducted in a clear, small-volume 384-well plate (# 784101, Greiner Bio-one). Fluorescence (535/585 nm) was measured after 45 min at using the POLARstar Omega (BMG LABTECH). Samples were normalized to protein content. Data were fitted (dashed line) using linear regression model in GraphPad Prism 5.0. Trolox standard curve was used to convert RSV results into Trolox equivalents and data were fitted (dashed line) using linear regression model.

### Intracellular pyruvate quantification

Intracellular pyruvate concentration was determined using the Pyruvate Assay Kit (# 700470, Cayman Chemical). NHEK cells were seeded in 150 cm^2^ cell culture flasks (Corning) and treated with 50 μM RSV or vehicle for 16 hours. Sample preparation was done according to the manufacturer’s instruction. Fluorescence (530/585 nm) was measured after 20 minutes using the POLARstar Omega (BMG LABTECH) at 37°C. Samples were normalized to protein content. Data were analysed using GraphPad Prism 5.0.

### Intracellular lactate quantification

Lactate content was quantified with the Lactate Assay Kit (# 700510, Cayman Chemical). NHEK cells were seeded in a 150 cm2 (Corning) and treated with 50 μM RSV or vehicle for 16 hours. Sample preparation was done according to the manufacturer’s instructions. Fluorescence (530/585 nm) was measured using the POLARstar Omega (BMG LABTECH) at 37°C. Samples were normalized to protein content. Data were analysed using GraphPad Prism 5.0.

### Analyses of intracellular ADP and ATP

The ADP Colorimetric/Fluorometric Assay Kit (K355-100) and ATP Colorimetric/Fluorometric Assay Kit (K354-100, both Biovision, BioCat) were used to quantify intracellular ADP and ATP content. Briefly, NHEK cells were seeded into 150 cm cell culture flasks (Corning) and treated with 50 μM RSV or vehicle for 16 hours. Flasks were washed once with ice cold PBS prior to harvest using a dispenser (TPP). Cell suspensions were centrifuged at 1,000 g for 5 minutes at 4°C and resuspended in ice-cold extraction buffer, aliquoted and stored at -20°C until usage. The assay was miniaturized to 10% of the initial volume and conducted in a clear, small-volume 384-well plate (# 784101, Greiner Bio-one) according to the manufacturer’s protocol. Optical density was measured after 45 minutes at 570 nm using the POLARstar Omega (BMG LABTECH). Samples were normalized to protein content. Data were analysed using GraphPad Prism 5.0.

### Intracellular NAD^+^ and NADH quantification

Intracellular NAD^+^ and NADH content was analysed using the colorimetric NAD/NADH Quantitation Kit (Biovision, BioCat) according to the manufacturer’s instruction. Briefly, NHEK cells were seeded into 150 cm cell culture flasks (Corning) and treated with 50 μM RSV or vehicle for 16 hours. Flasks were washed once with ice cold PBS prior to harvest using a dispenser (TPP). Cell suspensions were centrifuged at 1,000 g for 5 minutes at 4°C and resuspended in ice-cold extraction buffer. Afterwards, cells were lysed by two freeze-thaw-cycles, followed by intensive vortexing and centrifugation at 20,800 g for 15 minutes at 4°C. Part of the cell lysate was incubated at 60°C for 30 minutes to generate NAD^+^. Cycling buffer, enzyme mix and developer were added according to the manufacturer’s protocol. Optical density was measured after 30 minutes at 660 nm using the POLARstar Omega (BMG LABTECH). Samples were normalized to protein content. Data were analysed using GraphPad Prism 5.0.

### Analyses of intracellular NADP^+^ and NADPH

The NADP/NADPH-Glo Assay Kit (G9081, Promega) was used to quantify the intracellular NADP^+^ and NADPH content. NHEK cells were seeded into a 12-well plate and treated with 50 μM RSV or vehicle for 16 hours. The following day, cells were washed once in PBS, 60 μl PBS/well were added and cells were lysed in 60 μl base solution with 1 % dodecyl(trimethyl)azanium bromide (DTAB, D8638, Sigma Aldrich). Afterwards, 50 μl lysate were transferred into a clear 96-well plate to measure NADP^+^ and NADPH individually according to the manufacturer’s protocol.

Finally, 30 λl sample were transferred to a white-walled clear-bottom 384-well plate (# 781098, Greiner Bio-one) and 30 μl of NADP/NADPH-Glo Detection Reagent were added. Luminescence was measured after 30 minutes using the POLARstar Omega (BMG LABTECH). Samples were normalized to protein content. Data were analysed using GraphPad Prism 5.0.

### Analyses of intracellular reduced and oxidized glutathione

Intracellular reduced (GSH) and oxidized (GSSG) glutathione were quantified using the GSH/GSSG-Glo Assay Kit (V6611, Promega). NHEK cells were seeded in a 96-well plate (TPP) with a density of 30,000 cells/well. The following day, cells were treated with 50 μM RSV or vehicle for 16 hours. The assay was miniaturized to 25 % of the initial volume and conducted according to the manufacturer’s protocol. In brief, cell culture medium was removed, 12.5 μl/well Total Glutathione Lysis Reagent or Oxidized Glutathione Lysis Reagent were added and incubated for 5 min at RT while shaking. Afterwards, 12.5 μl/well Luciferin Generation Reagent were added and incubated at RT for 30 minutes. Samples and standards were transferred to a white 96-well plate (# 655083, Greiner Bio-one) and 25 μl Luciferin Detection Reagent were added. Luminescence was measured after 15 minutes using the POLARstar Omega (BMG LABTECH). Samples were normalized to protein content. Data were analysed using GraphPad Prism 5.0.

### Oxygen consumption assay

Analyses of oxygen consumption was conducted by using the cell impermeable, oxygen-sensing fluorophore MitoXpress Xtra (Luxcel Biosciences). NHEK cells were seeded in a 96-well plate (TPP, Biochrom) with a density of 25,000 cells/well. The day after, cells were treated with 50 μM RSV or vehicle for 16 hours and afterwards equilibrated for 20 minutes under CO_2_-free conditions at 37°C. After aspirating cell culture medium, 62.5 nM/well MitoXpress Xtra diluted in KGM (LONZA) was added. The plate was incubated for 10 minutes under CO_2_-free conditions at 37°C. Compounds were added as indicated and each well was sealed with 100 μl HS Mineral Oil (Luxcel Biosciences). Dual-read time-resolved fluorescence (TR-F) was measured (380/650 nm) with the POLARstar Omega (BMG LABTECH) at 37°C. Lifetime, dual-delay and gate times were calculated and set according to the manufacturer’s instructions. Data were analysed using GraphPad Prism 5.0.

### Fluorescence microscopy

NHEK cells were seeded in X-well tissue culture chambers (Sarstedt, Nurnbrecht, Germany) at a density of 20,000 cells/well. One day later, adherent cells were treated for 16 hours with the indicated compounds. Visualization of MAP1LC3A, Actin, Nrf2, Keap1, GCLC and GSR was done as recently described in Weidner, et al. ^43^. In brief, cells were washed with PBS (Sigma Aldrich) and fixed in 4 % formaldehyde/PBS (Sigma Aldrich) for 15 minutes at RT. Cells were washed with PBS (Sigma Aldrich), permeabilised in 0.3 % Triton X-100/PBS (PBS-Tx) for 10 minutes at RT, washed with PBS and afterwards blocked in 5 % goat serum (Sigma Aldrich) in PBS-Tx for 60 minutes at RT. Afterwards, cells were incubated with primary antibodies at 4°C overnight. Primary antibodies and dilutions were as follows: MAP1LC3A (1:100; AP1801d-ev-AB, Biovision, BioCat), Nrf2 (1:100; sc-13032X, Santa Cruz; 1:100), Keap1 (1:100, sc-365626, Santa Cruz), GCLC (1:100, ab41463, Abcam) and GSR (1:100, sc-133159, Santa Cruz). Subsequently, cells were washed with PBS-Tx and stained with anti-mouse IgG (H+L) F(ab')2 fragment Alexa Fluor 555 Conjugate (# 4409, Cell Signaling Technology) and anti-rabbit IgG (H+L), F(ab')2 fragment Alexa Fluor 488 Conjugate (# 4412, Cell Signaling Technology) diluted (1:1,000) in 1 % BSA/PBS-Tx for 1 h at RT in the dark. Subsequently, cells were washed with PBS for 5 minutes, and the MAP1LC3A stained cells were counterstained with F-actin probe Texas Red-X Phalloidin (1:100, T7471, Life Technologies) for 20 minutes at RT in the dark. Cells were washed with PBS, the chamber was removed from the slide, samples were counterstained with ProLong Gold Antifade Mountant (with DAPI) solution (Life technologies), a cover slip applied (# 235503704, Duran Group, Wertheim/Main, Germany) and incubated at RT for 24 hours. Fluorescence microscope imaging was performed on the LSM700 (Zeiss, Jena, Germany).

### Redox potential, redox state and redox environment

To calculate the half-cell reduction potential of a selected redox couple the Nernst equation (Eq. 2) was used and transformed to match the experimental conditions (T = 310K = 37°C, pH 7) (Eq. 3).

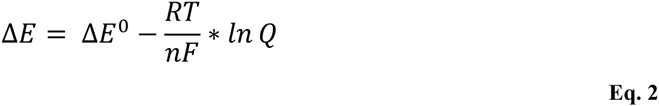

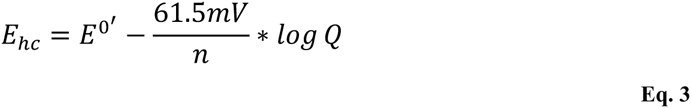

R is the gas constant (8.314 J K^-1^ mol^-1^), T the temperature in Kelvin, n the number of electrons exchanged, F the Faraday constant (9.6485*10^4^ C mol^-1^) and Q the mass action expression. As the standard reduction potential (E^0^, E^0^) is pH dependent, an adjustment is necessary:

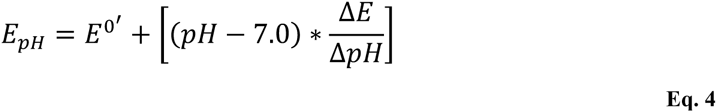

E_pH_ represents the half-cell reduction potential at a given pH, while ΔE/ΔpH is dependent on the number of electrons and protons involved. Finally, the half-cell reduction potential of a selected redox couple can be calculated according to (Eq. 5).

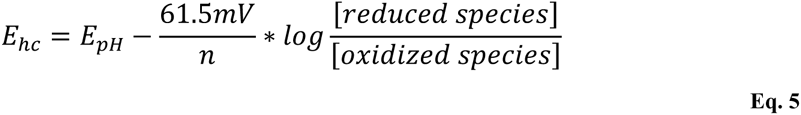

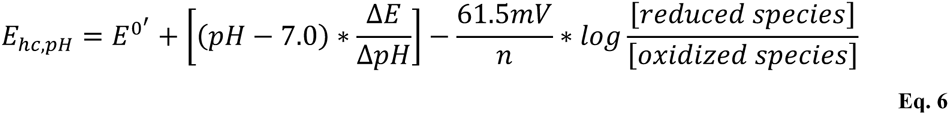

The half-cell reduction potential can be calculated for the following metabolites (redox couples):

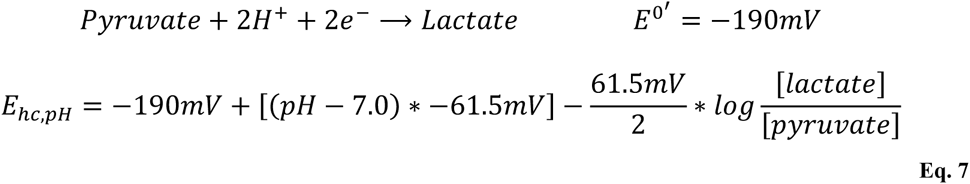

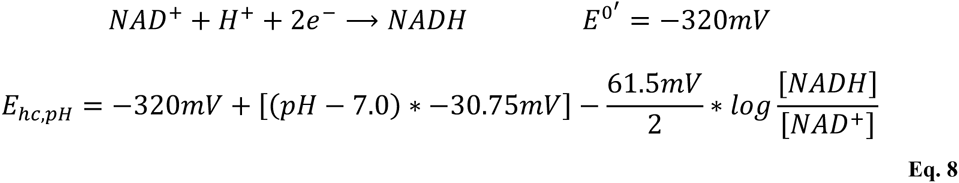

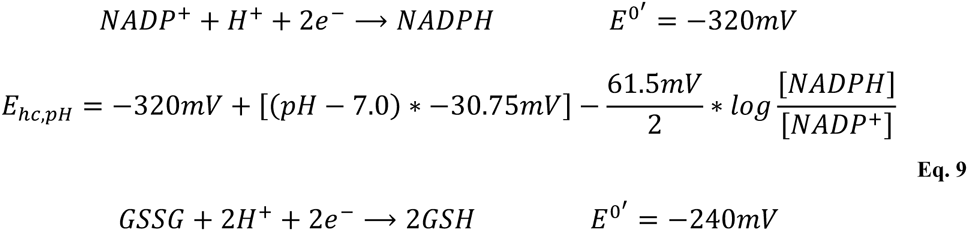

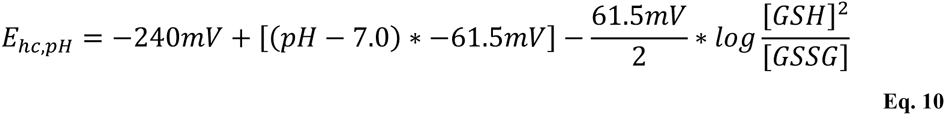

Finally the redox environment is calculated using the redox couple 2GSH/GSSG alone (Figure 5a and Table S4) or redox couples lactate/pyruvate, NADH/NAD^+^ and 2GSH/GSSG (Fig. S8f and Table S4), respectively.

To relate the reduced redox environment to a future oxidative challenge, we did a thought experiment focusing on oxidation of ethanol to acetaldehyde and simultaneous ROS generation ^7-9^:

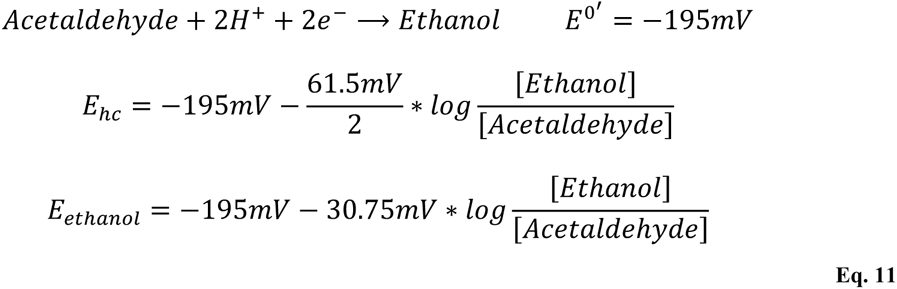

In order to estimate the amount of ethanol caused ROS generation, which can be quenched by RSV pre-treatment, we made the following assumptions:

- pH 7,
- T = 37°C,
- 2GSH/GSSG redox couple is the driving force of the redox environment and
- Concentrations of GSH and GSSG as measured at pH 7.0026 (vehicle) and 7.0632 (RSV), respectively.

At equilibrium, change of the half-cell reduction potentials of both couples equal 0 and can be transformed appropriately:

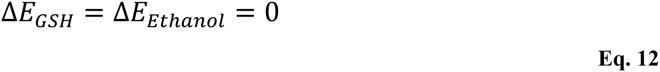

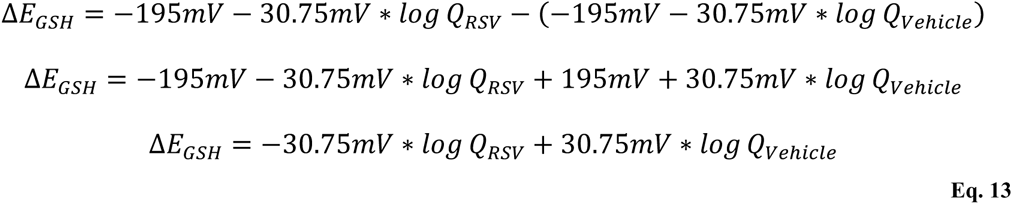

According to Sarkola, et al. ^44^ the formation of acetaldehyde in humans is dependent on the amount of ethanol. The mean slope was determined as follows:

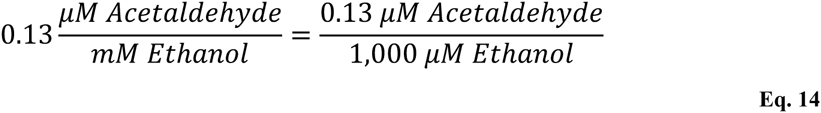

We can use this information to calculate Q_Vehicle_:

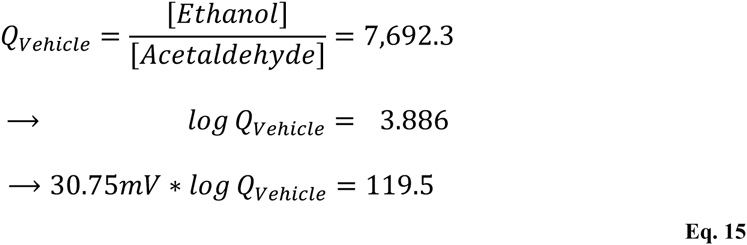

Finally we can calculate Q_RSV_ accordingly:

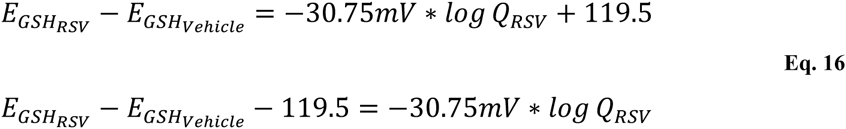

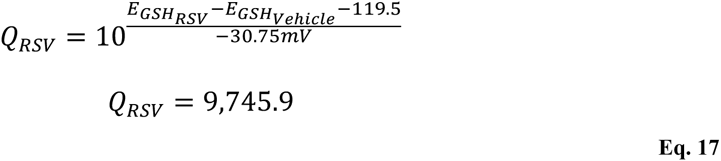

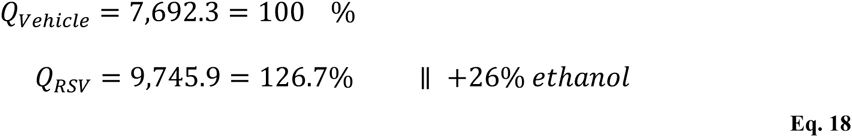

Thus, we can estimate that owing to RSV pre-treatment and the resulting shift in redox environment, NHEKs are likely to tolerate roughly 26 % more ethanol than vehicle pre-treated cells. Notably, NHEK cells did not tolerate higher doses of ethanol without serious reduction of viability. Consequently, experimental evidence for this gedankenexperiment can finally not be provided.

We further challenged RSV pre-treated NHEKs with HNE, a potent thiol scavenger ^39^,40,^45^. As 1 molecule GSH is needed to detoxify 1 molecule HNE, we can roughly estimate the amount of additional HNE quenched by increased cellular concentrations of GSH after RSV pre-treatment. While RSV pre-treatment has still a protective effect at 15 μM HNE, this protection is abrogated at 25 μM HNE. Corresponding to the increased GSH content (7 μmol per g protein, roughly 2 μM) due to pre-treatment with RSV, the pre-conditioning effect is depleted at HNE concentrations ≥ 25 μM. Nevertheless, this is just an approximation, as the GSH pool regulates the reduction state of many biological molecules and because HNE can efficiently react with many intracellular thiol groups, lipids and proteins present in the biological system.

### Statistical analyses

Data are expressed as mean ± standard error (s.e.m.), if not otherwise denoted. Statistical tests were performed in GraphPad Prism 5.0. For comparison of two groups statistical significance was examined by unpaired two-tailed Student’s t-test. One-way ANOVA with subsequent Dunnett’s post-test was used for multiple comparisons. A *P* value ≤ 0.05 was defined as statistically significant.

## Acknowledgements

We thank Gerald Rimbach, Sophia Bauch, Stefanie Becker and Chung-Ting Han for valuable discussion and for support.

## Author contributions statement

S.S. conceived, designed and supervised the study. A.P., A.G., S.C., S.J.W., L.L., C.W., M.K. G.J., S.L., L.J.W. and S.S. designed the experiments. A.P., A.G., S.C., S.J.W., L.L., M.R., and L.F. performed the experiments. A.P., A.G., S.C., S.J.W., L.L., L.F. and S.S. analysed the data. G.J., S.L., L.J.W. and S.S. provided tools. A.P. and S.S. wrote the paper with input from M.K. and all other authors.

## Additional Information Accession codes

The accession number for all genome-wide gene expression data used in this study is GEO: GSE72119.

